# Gene expression patterns associated with fin shape differ between two lamprologine cichlids

**DOI:** 10.1101/2022.06.02.494591

**Authors:** Ehsan Pashay Ahi, Florian Richter, Kristina M. Sefc

## Abstract

Comparing gene regulatory patterns between seemingly similar phenotypic traits can provide important insights on the molecular mechanisms underlying the evolution of those traits. In this study, we investigate the molecular basis of the formation of a spade-shaped caudal fin, which is a rare phenotype among teleost fish characterized by an elongated medial region of the fin. We examined the expression patterns of candidate fin-shape genes in the spade-shaped caudal fin of the related species *Lamprologus tigripictilis*, an East African cichlid in the tribe Lamprologini. The candidate gene set consisted of a previously identified gene regulatory network (GRN) associated with the elongation of fin regions in another Lamprologini cichlid species and further genes selected on the basis of co-expression data and transcription factor prediction. Unexpectedly, the anatomical features of elongated fin rays differed and gene expression patterns associated with fin elongation were only weakly conserved between the two related species. We report 20 genes and transcription factors (including *angptl5, cd63, csrp1a, cx43, esco2, gbf1* and *rbpj*), whose expression levels differed between the elongated and the short caudal fin regions of *L. tigripictilis*, and which are therefore candidates for the regulation of the spade-like fin shape.

## Introduction

The shape of the fins of ray-finned fishes is determined predominantly by variation in the length of the individual fin rays. In many species, one or more of the fins are adorned by elongated filaments, which are formed by accelerated or prolonged growth of these fin regions during larval or later development. Less conspicuous shapes, such as spade- of fork-shaped caudal fins, are likewise the product of ray length variation within fins. Importantly, fin shapes can regenerate completely after damage or experimental amputation throughout the lifetime of the fish, suggesting that it is under rather strict genetic control and that positional memory orchestrates the molecular factors necessary for fin regeneration (Rabinowitz et al., 2017).

Most of the ground-laying work on the genetic control of fin growth and shape has concentrated on zebrafish (Pfefferli & Jaźwińska, 2015), but fish species may be expected to differ with respect to the molecular and anatomical basis of fin shape formation. The fin rays of teleost fish are segmented, and both ontogenetic and regenerative growth involves the addition of bony segments at their distal ends. Ray length is therefore determined by either or both of the length and the number of individual segments. At the molecular level, this involves components of various developmental pathways, such as WNT, FGF, Hedgehog and retinoic acid (RA) signalling, as well as epigenetic, skeletogenic and structural remodelling factors (Iovine, 2007; Yoshinari et al., 2009; Wehner & Weidinger, 2015; Sehring & Weidinger, 2020).

We have previously conducted a study of the anatomy and the molecular mechanisms underlying the ornamental fin shape of the cichlid fish *Neolamprologus brichardi*, an endemic of Lake Tanganyika in East Africa (Ahi et al., 2017; Ahi & Sefc, 2018). In this species, the distal tips of the dorsal and anal fins as well as the dorsal and ventral tips of the fork-shaped caudal fin are conspicuously elongated. This shape allowed us to compare expression levels of candidate genes between elongated and non-elongated (short) regions within the same fins, and we did so by using intact as well as regenerating fin tissue sampled from adult fish. The observed gene expression patterns and correlations led to the proposition of a gene regulatory network (GRN) involved in the formation of the fin phenotype, hitherto referred to as *N*.*b*.-GRN. Members of the *N*.*b*.-GRN include genes reported to be involved in fin ray segmentation, angiogenesis or neurogenesis such as *cx43, mmp9, angptl5, angptl7, dpysl5a, csrp1a* and *cd63* (Iovine et al., 2005; Monaghan et al., 2006; Nakatani et al., 2007; Sims et al., 2009; Ma et al., 2012; Ton & Iovine, 2012, 2013; Kang et al., 2015; Hagedorn et al., 2016), and several potential upstream regulators for this gene network including *egr2, foxc1, foxd3, foxp1, irf8* and *myc* (Ahi & Sefc, 2018). Among the latter, *myc, irf8* and *foxd3* were already indicated in fin regeneration studies of other teleost fish (Christen et al., 2010; Li et al., 2012; Kang et al., 2015; Hasegawa et al., 2017; Huang & Chen, 2017). We predicted *foxd3* as the key upstream regulator of the gene network in *N. brichardi*, since it consistently displayed significant expression correlation with all members of the network genes (Ahi & Sefc, 2018).

We next investigated whether members of the *N*.*b*.-GRN were also implicated in similar elongations of the dorsal and anal fins in another East African cichlid species, *Steatocranus casuarius*, a member of Steatocranini tribe which is a sister lineage to the Lake Tanganyika cichlids (Ahi et al., 2019). The caudal fin of this species does not exhibit any pronounced elongations and was therefore not studied. Only a subset of the tested genes showed the expected expression level differences between short and elongated regions of anal and dorsal fins, indicating that the molecular mechanisms controlling fin elongation differ between cichlid fish. Gene expression patterns in comparisons of elongated and short fin regions also differed within *S. casuarius*; for instance, several genes showed significant differences in the conspicuously elongated dorsal fin but not in the anal fin, where the elongation is less pronounced (Ahi et al. 2019).

The divergence in the molecular mechanisms of fin elongations between the two species was accompanied by anatomical differences. In *N. brichardi*, fin ray segments in the elongated fin regions were shorter or, in case of the caudal fin, of the same length as the fin ray segments in the short fin regions, suggesting that the elongation must be due to a larger number of segments in the elongated rays (Ahi et al., 2017). In *S. casuarius*, in contrast, segments of the elongated rays were longer than the fin ray segments in the short fin region (Ahi et al., 2019). The number of segments per ray was not determined, but as the segment length difference between short and elongated rays was not pronounced enough to explain the fin shape, the rays in the elongated fin regions of *S. casuarius* probably consist of more and longer segments.

The two species, *N. brichardi* and *S. casuarius*, represent two distantly related cichlids with convergent shape of the dorsal and anal fins (Ahi et al., 2019), with a divergence time of about 14 million years between their respected tribes, the Lamprologini (*N. brichardi*) and the Steatocranini (*S. casuarius*; Irisarri et al. 2018). In the present study, we examine the caudal fin of *Lamprologus tigripictilis*, which is reverse to the shape of the caudal fin in *N. brichardi* – specifically, a spade-shaped caudal fin, in which the medial region is elongated compared to the short dorsal and ventral regions of the fin. *Lamprologus tigripictilis* is more closely related to *N. brichardi*, as the radiation of the Lamprologini, to which both species belong, started at about 6 million years ago (Irisarri et al., 2018). This sets a considerably lower upper limit on the divergence time between these two species than that between *N. brichardi* and *S. casuarius*. If mechanisms of fin ray elongation are shared between the two lamprologine cichlids, we expect that many of the *N*.*b*.-GRN member genes, that have been identified in *N. brichardi*, will show corresponding gene expression differences between the elongated and short caudal fin regions in *L. tigripictilis*. We first examined the caudal fin expression patterns of 16 members of the *N*.*b*.-GRN by comparing gene expression levels in short and elongated regions using qPCR, and then expanded the set of candidate genes based on co-expression data and transcription factor prediction in order to identify additional members of the *N*.*b*.-GRN operating in *L. tigripictilis*. We also measured the length of the fin ray segments in the short and elongated fin rays in order to assess the anatomical convergence of fin elongation in *L. tigripictilis* with the other two cichlid species.

## Methods

### Fin sampling for RNA isolation and cDNA synthesis

We used six captive bred adult males of *Lamprologus tigripictilis* (total length 7-10cm). Prior to taking the fin biopsies, fish were anesthetized using 0.04 gram of MS-222 per liter of water. Then, the caudal fin was cut in front of the first ray bifurcation (branching) under a stereomicroscope (red dashed lines in Fig. 1A, B). Next, three separate tissue samples were dissected from the severed fin: one comprised the most dorsal, branched fin ray and represented the dorsal short fin region (dS in Fig. 1), one comprised the most ventral, branched fin ray represented the ventral short fin region (vS in Fig. 1) and one comprised the medial fin ray and represented the medial elongated fin region (mL in Fig. 1). Each sample comprised the two branches of the selected fin ray. The tissue samples were stored frozen in RNAlater (Qiagen) until RNA isolation.

**Figure 1:**
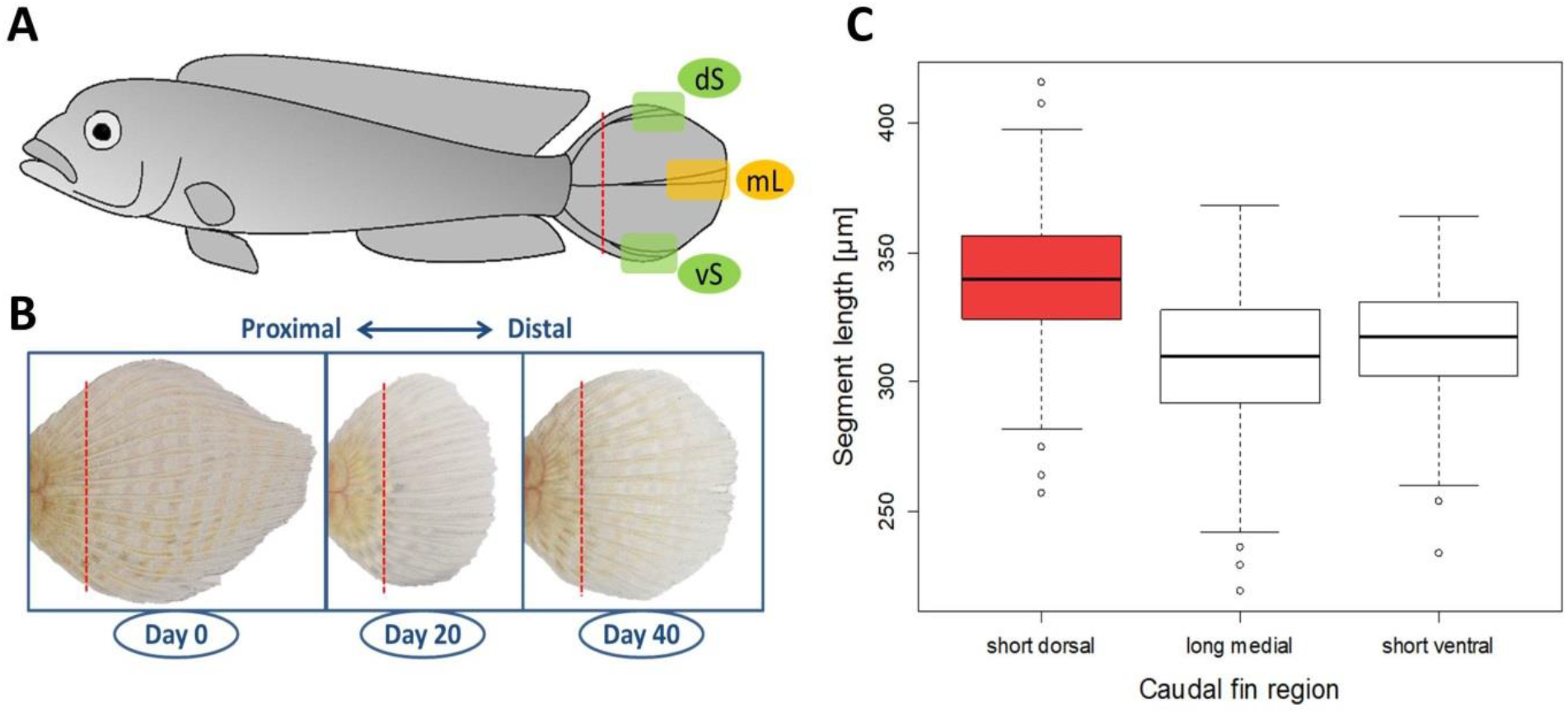
The caudal fin of *Lamprologus tigripictilis*. (A) Caudal fins were amputated along the dashed red line, and tissue samples representing the short dorsal (dS), the short ventral (vS) and the elongated medial (mL) regions of the dorsal fin were taken as indicated. (B) Biopsies were taken from the original fin tissue (day 0, representing stage 0), and from regenerating tissue at day 20 (representing stage 1) and day 40 (representing stage 2) after the preceding amputation. (C) Length of fin ray segments in the three regions of the caudal fin. Different box colours indicate significant differences in segment length.

Gene expression was examined in biopsies of the original fin tissue (stage 0) to study gene activity patterns associated with the maintenance of the phenotype, and twice during regeneration, including a biopsy at day 20 after the stage0 biopsy when the round shape of the fin started to appear (stage 1), and another biopsy at day 40 after the stage1 biopsy, when the fin had become distinctly rounded (stage 2; Fig. 1B). RNA isolation and cDNA synthesis was performed as described in our previous studies (Ahi et al., 2017). Anesthesia and fin biopsies were performed under permit number BMWFW-66.007/0028-WF/V/3b/2017 issued by the Federal Ministry of Science, Research and Economy of Austria (BMWFW). All methods were performed in accordance with the relevant guidelines and regulations of BMWFW.

### Candidate target and reference gene selection

The selection and analysis of candidate target genes and transcription factors (TF) was performed in three steps (see Ahi et al. 2015). First, we tested 16 genes of the *N*.*b*.-GRN described in the introduction (Ahi & Sefc, 2018). These genes were *angptl5, angptl7, anxa2a, c1qtnl5a, cd63, csrp1a, cx43, dpysl5a, gnao1a, kif5a, mmp9, pfkpa, sema3d, txn, wnt10a* and *wnt5b*. Out of the first set of candidate genes, we picked the ones with the strongest expression differences between the elongated and the short fin regions (*angptl5, cd63, csrp1a*; see results) to search for genes co-expressed with all the three genes (top overlapping coexpressed genes) in the zebrafish co-expression database, COXPRESdb (http://coxpresdb.jp/) version 6.0 (Obayashi & Kinoshita, 2011). To attain a high degree of reliability, we filtered the genes co-expressed with each of the three genes by setting the Supportability score to a minimum of 1 (as described by Obayashi & Kinoshita, 2011) (Supplementary data 1). This step identified eight additional candidate genes (see results). Finally, we selected eight of the above 24 candidate genes, namely those with strongly increased expression in the elongated fin region (see results), for TF prediction. In order to predict the potential upstream regulators for these genes, we performed motif enrichment on 4 kb upstream sequences (promoter and 5’-UTR) of these genes as previously described by (Lecaudey et al., 2019) using the annotated genome of the Nile tilapia (Flicek et al., 2013) and two algorithms: MEME (Bailey et al., 2009) and XXmotif (Luehr et al., 2012). The motifs that were present in the promoters of at least half of these genes were compared to position weight matrices (PWMs) from the TRANSFAC database (Matys et al., 2003) using STAMP (Mahony & Benos, 2007) to identify matching transcription factor (TF) binding sites (Supplementary data 1).

To identify stable reference genes, we selected 8 candidate reference genes with abundant expression in a range of tissues, which have already been investigated as reference genes in fins or other tissues containing skeletal structures or/and epidermis in fish (Table 1). Candidate reference genes were ranked according to expression stability by three different algorithms, BestKeeper (Pfaffl et al., 2004), NormFinder (Andersen et al., 2004) and geNorm (Vandesompele et al., 2002). The standard deviation (SD) based on Cq values of the fin regions was calculated by BestKeeper to determine the expression variation for each reference gene. In addition to ranking, BestKeeper also determines the stability of reference genes through a correlation calculation or BestKeeper index (r). GeNorm calculates mean pairwise variation between each gene and other candidates (the expression stability or *M* value) in a stepwise manner and NormFinder identifies the most stable genes (lowest expression stability values) based on analysis of inter- and intra-group variation in expression levels variations (Ahi et al., 2013). All three algorithms ranked *rps18* and *actb1* as the two most stable candidate reference genes (Table 1). Based on these results, we used the geometric mean of the expression of *actb1* and *rps18* for normalization of relative gene expression of candidate target genes.

**Table 1.** Ranking and statistical analyses of reference genes in the caudal fin of *L. tigripictilis* using three different algorithms. Abbreviations: SD = Standard deviation, r = Pearson correlation coefficient, SV = stability value, M = M value of stability.

| BestKeeper |  |  |  | geNorm |  | NormFinder |  |
| --- | --- | --- | --- | --- | --- | --- | --- |
| Ranking | SD | Ranking | r | Ranking | M | Ranking | SV |
| *rps18* | 0.975 | *rps18* | 0.732 | *actb1* | 0.530 | *actb1* | 0.183 |
| *actb1* | 0.973 | *actb1* | 0.903 | *rps18* | 0.548 | *rps18* | 0.229 |
| *hsp90a* | 0.958 | *hsp90a* | 0.946 | *hsp90a* | 0.562 | *rps11* | 0.246 |
| *rps11* | 0.942 | *rps11* | 0.992 | *rps11* | 0.604 | *hsp90a* | 0.254 |
| *tbp* | 0.902 | *hprt1* | 1.013 | *hprt1* | 0.685 | *tbp* | 0.377 |
| *hprt1* | 0.889 | *tbp* | 1.061 | *tbp* | 0.690 | *hprt1* | 0.418 |
| *elf1a* | 0.798 | *elf1a* | 1.197 | *elf1a* | 0.904 | *elf1a* | 0.512 |
| *gapdh* | 0.752 | *gapdh* | 1.212 | *gapdh* | 0.976 | *gapdh* | 0.540 |

### Primer design and real-time qPCR

In order to design qPCR primers, we aligned the orthologues of each gene from different African cichlid tribes including one species of Tilapiini (*Oreochromis niloticus*), one species of Lamprologini (*N. brichardi*) and one species of Haplochromini (*Astatotilapia burtoni*) (Brawand et al., 2014; Santos et al., 2016; Singh et al., 2017). The 1-to-1 orthologues were confirmed by blasting zebrafish mRNA REfSeq IDs against *N. brichardi* transcriptome in NCBI and cross-checking the top hits returned by BLAST in the Ensembl database for zebrafish and *O. niloticus* orthologues (http://www.ensembl.org). Next we used the aligned sequences to identify conserved regions across the species (using CLC Genomic Workbench, CLC Bio, Denmark) and at the exon/exon boundaries (using annotated genome of *O. niloticus* in the Ensembl database. Primers with short amplicon sizes (<250 bp) were designed using Primer Express 3.0 (Applied Biosystems, CA, USA) and OligoAnalyzer 3.1 (Integrated DNA Technology) (Supplementary data 1), as previously described (Ahi et al., 2020).

The qPCR was conducted using Maxima SYBR Green/ROX qPCR Master Mix (2X) by following the manufacturer’s instruction (Thermo Fisher Scientific, St Leon-Rot, Germany) in 96 well-PCR plates on an ABI 7500 real-time PCR System (Applied Biosystems). The experimental set-up per run followed the preferred sample maximization method (Hellemans et al., 2007). The primer efficiency analyses in LinRegPCR v11.0 (http://LinRegPCR.nl) (Ramakers et al., 2003) were conducted as described in our previous study (Ahi et al., 2017).

### Analysis of qPCR data

The geometric mean of the Cq values (Vandesompele et al., 2002) of the two reference genes Cq _reference_, was used to normalize Cq values of target genes in each sample (ΔCq _target_ = Cq _target_ – Cq _reference_). In order to calculate ΔΔCq values, we randomly selected one biological replicate of the dorsal region of the caudal fin (stage 0) and subtracted the ΔCq from the calibrator ΔCq value (ΔCq _target_ – ΔCq _calibrator_). Relative expression quantities (RQ values) were calculated as 2^−ΔΔCq^ (Pfaffl, 2001).

RQ values were log-transformed for statistical analyses. For each target gene, differences in expression levels between dorsal (short) and medial (elongated), ventral (short) and medial (elongated) as well as dorsal (short) and ventral (short) regions of the caudal fin were tested in linear mixed models with log(RQ) as dependent variable, fin region as fixed factor and developmental stage nested within biological replicate (fish) as grouping factor (Supplementary data 2). We conducted paired t-test to calculated p-values and to account for multiple testing (N = 111 comparisons; 37 candidate genes times 3 fin region contrasts), p-values for the effect of length were corrected using the Benjamini-Hochberg procedure (Benjamini & Hochberg, 1995). For some genes, we also report results of analyses which included only stage 1 and stage 2 data (i.e. the regenerating tissue but not the intact tissue). The log-transformed RQ values were also used for calculation of pairwise Pearson correlation coefficients (*r*) and associated p-values.

### Fin ray segment length measurements

To measure the length of fin ray segments in elongated medial and the short dorsal and ventral regions of the caudal fin, biopsies were taken from six adult males as described above for RNA sampling and stained with alizarin red. We modified the acid-free double staining protocol described by (Walker & Kimmel, 2007) and used 10% KOH in the clearing phase, increased the duration of the staining phase to 4 days and the duration of the clearing phase to 15 days. Using a Keyence VHX-5000 Digital Microscope, we measured the lengths of the 10 most distal, complete segments on one branch per each of two dorsal (short), two ventral (short) and four medial (elongated) fin rays. That is, measurements were taken from eight different rays, and not from branches pertaining to the same ray. The selected fin rays were the two most dorsal, branched rays, the two most ventral, branched rays (we did not use the most dorsal and the most ventral fin ray, as these were only rudimentary developed in some individuals) and the four medial fin rays. The branch representing each fin ray was either selected randomly or by avoiding irregularities in the segmentation pattern, which sometimes occurred in one of the branches. We used a linear mixed model (R package lmerTest) to compare segment length between the three fin regions. To account for possible correlations within rays and individual fins, we included ‘fin ray’ nested in biological replicate (fish) as grouping factor.

## Results

### Morphological characterization of the caudal fin

The lengths of the fin ray segments (Fig. 1) did not differ significantly between the elongated, medial fin region (mean ± sd = 309.4 ± 25.5 µm) and the short, ventral fin region (mean ± sd = 315.1 ± 25.2 µm; est.= 5.9, t = 1.6, p = 012). In contrast, the individual fin ray segments were significantly longer in the short, dorsal fin region (mean ± sd = 339.6 ± 28.6 µm), both compared to the medial fin region (est. = 30.5, t = 8.2, p = 5.6 * 10^−10^) and compared to the ventral fin region (est. = 24.6, t = 5.7, p = 1.2 * 10^−6^).

Hence, comparisons between the medial and the ventral region of the caudal fin represent a contrast between long and short fin rays that differ in the number of segments per ray. In contrast, comparisons involving the dorsal fin region involve differences in both segment length (longer in dS) and number (fewer in dS), with a larger difference in the number of segments between dS and mL than between dS and vS.

### Expression analysis of candidate genes

Each of the three pairwise comparisons between fin regions represents a different phenotypic contrast (see above), and we therefore conducted three pairwise comparisons of candidate gene expression levels among the fin regions (as opposed to combining the data into a single ‘short’ versus ‘long’ comparison). In the following text, ‘expression in the medial elongated region of the caudal fin’ is abbreviated as ‘mL-expression’, ‘expression in the dorsal region’ is abbreviated as ‘dS-expression’ and ‘expression in the ventral region’ is abbreviated as ‘vS-expression’, and the expression levels are reported as ‘increased’ or ‘decreased’ in comparison to the other regions. We obtained similar results in analysis that included all three stages (intact fin and regeneration stages 1 and 2) and in analysis restricted to the two regeneration stages, and report the analysis of the full data unless noted otherwise. Table 2 summarizes the significant expression level differences detected among the tested candidate genes, and the complete statistical analyses are reported in the supplementary material (Supplementary data 2).

**Table 2:**
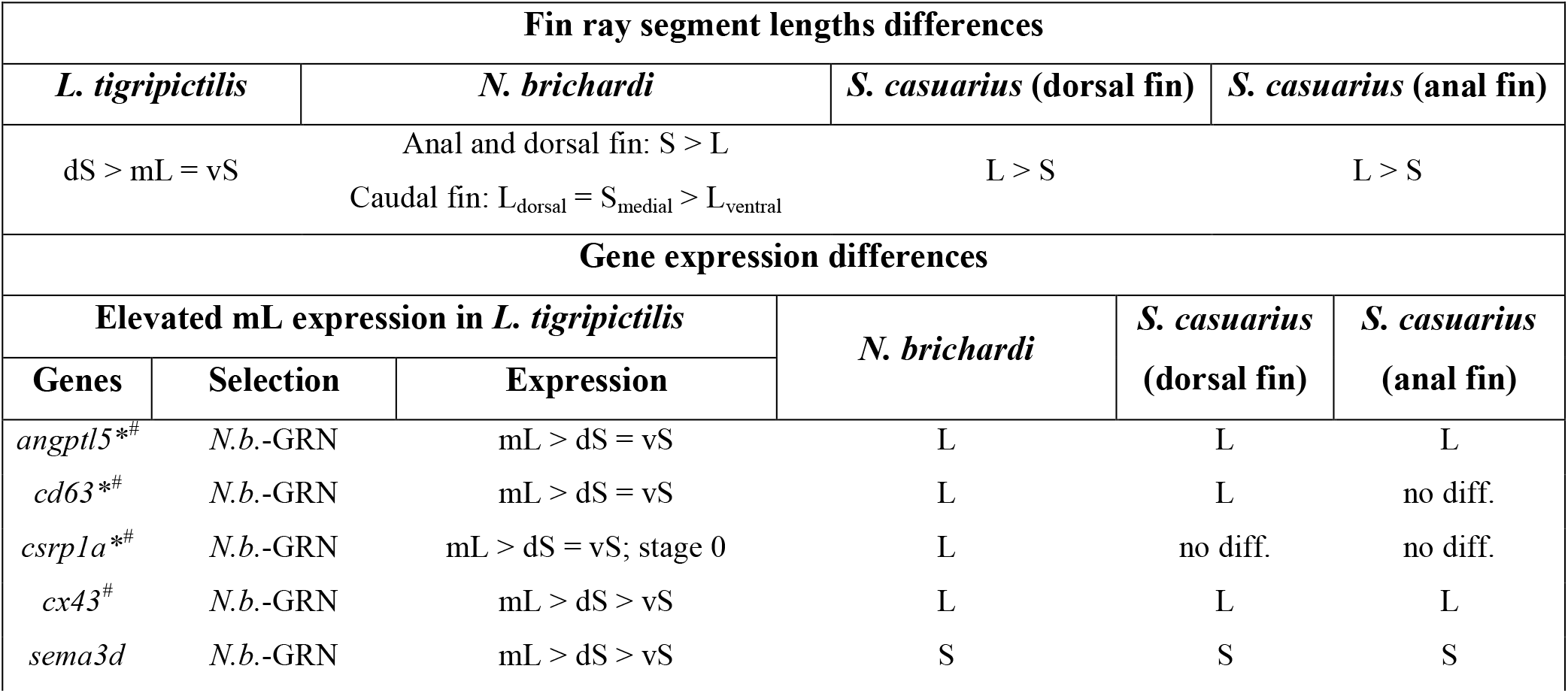
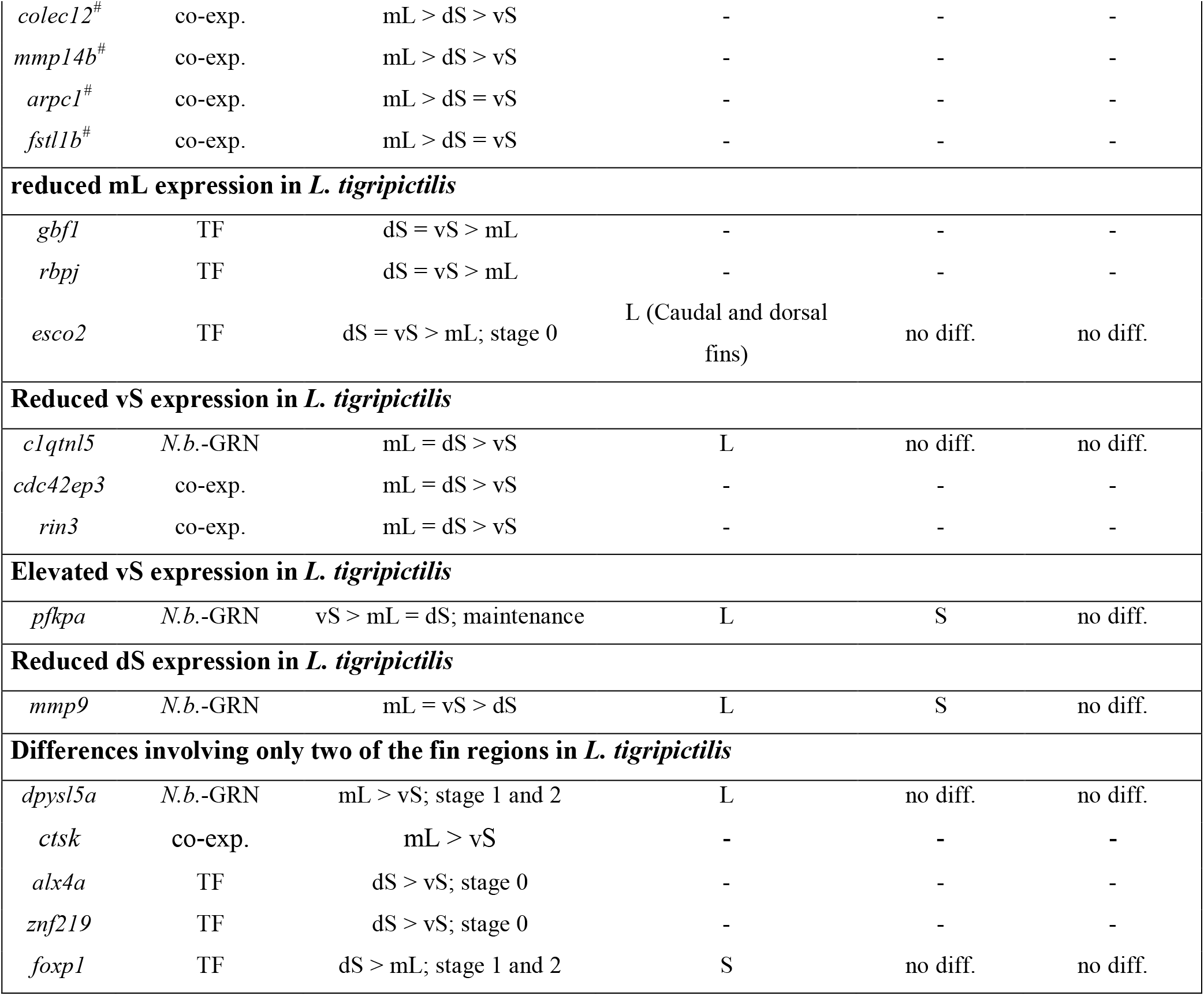
Summary of anatomical and gene expression patterns in the caudal fin regions of *L. tigripictilis*, compared to the patterns detected in two other cichlid species. mL, dS and vS are the medial long, dorsal short and ventral short regions of the caudal fin of *L. tigripictilis*, while L and S are the elongated and short regions of the fin types examined in *N. brichardi* and *S. casuarius*. In the summary of the segment length differences, dS > mL (for instance) indicates that segments in the dS region are longer than those in the mL region. In the summary of the gene expression differences, we report results for genes with significant expression level differences detected in *L. tigripictilis*, sorted by the detected pattern. ‘*N*.*b*.-GRN’ identifies candidate genes that are part of the gene regulatory network identified in *N. brichardi*; ‘co-exp.’ identifies candidate genes based on coexpression; ‘TF’ identifies the predicted transcription factors. Asterisks mark genes underlying the search for coexpressed candidate genes; hashes mark genes used for TF prediction. The gene expression pattern mL > dS = vS for *angptl5*, for instance, signifies that the expression level of the gene is significantly higher in mL compared to dS and to vS, whereas expression levels in dS and vS are not significantly different from each other. Unless developmental stages are indicated, the reported difference in gene expression levels was observed in the analysis including all stages. S and L stand for significantly elevated gene expression levels in short or long, respectively, regions of the fins of *N. brichardi* and *S. casuarius* detected in previous studies (Ahi et al., 2017, 2019; Ahi & Sefc, 2018). In *N. brichardi*, the expression patterns were largely consistent across the three fin types and also within the caudal fin (i.e., concerning the contrasts between the medial short region on the one hand and the dorsal and ventral elongated regions on the other hand); therefore, results could be summarized across fins unless indicated otherwise. ‘no diff.’ indicates that no significant expression L/S differences could be detected in *N. brichardi* or *S. casuarius*; when no pattern is reported for *N. brichardi* and *S. casuarius*, these genes were not tested in these species.

In the first step of our gene expression analysis, we examined the expression levels of 16 members of the *N*.*b*.-GRN. Among these, we detected increased mL-expression compared to both vS- and dS-expression for *angptl5, cd63, csrp1a, cx43* and *sema3d*, and increased mL-expression compared to vS-expression for *c1qtnf5* and *dpysl5a* (mainly in stage 1 of fin regeneration) (Fig. 2, Supplementary data 2). Additionally, dS-expression of *mmp9* was lower than mL- expression. We also detected significant expression level differences for *pfkpa* (higher mL- than dS-expression during regeneration).

**Figure 2:**
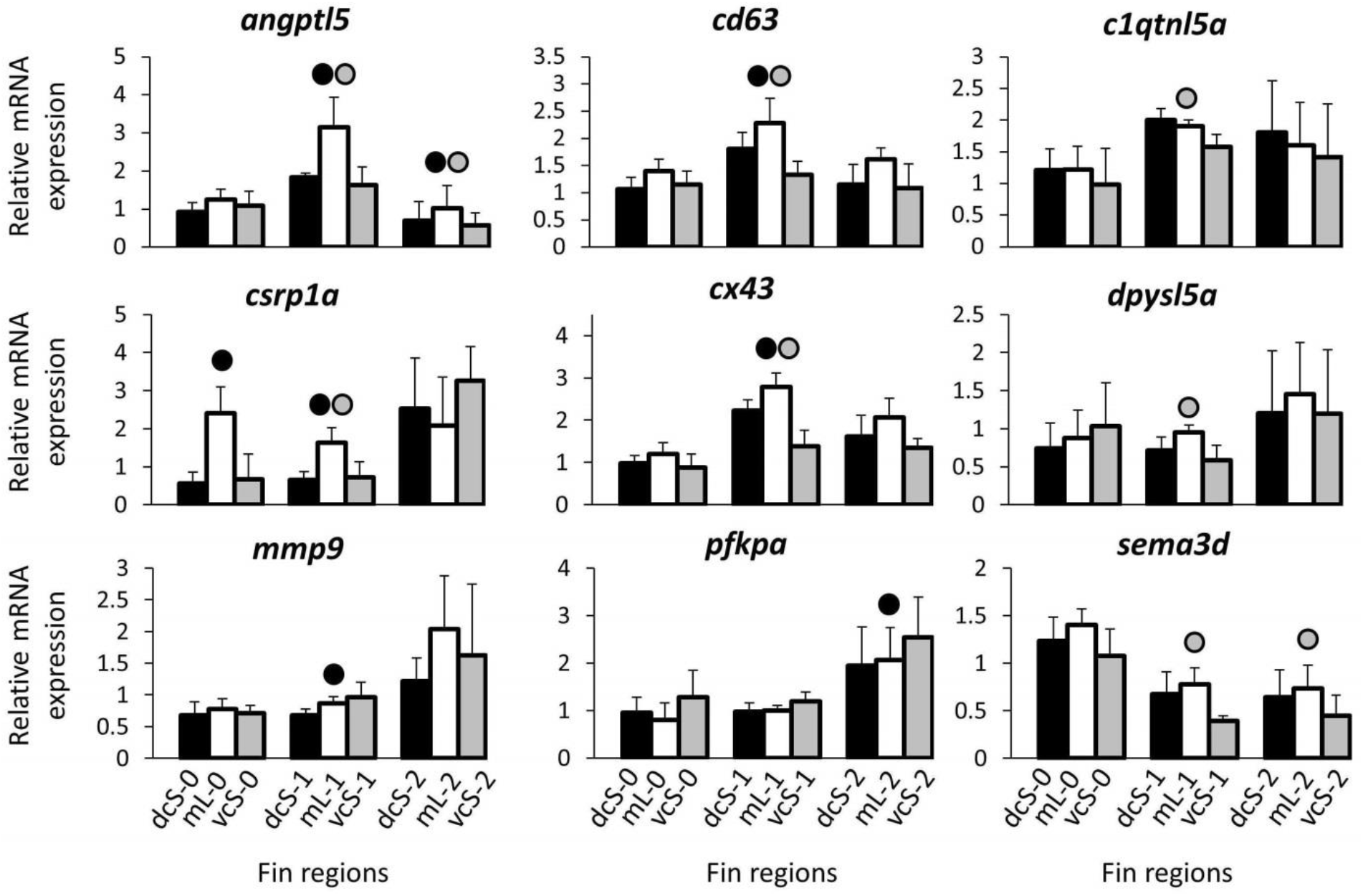
Expression levels of candidate genes selected based an already identified GRN in *N. brichardi*. Means and standard deviations of RQ in three biological replicates are shown for the elongated and short regions of the caudal in original (stage 0) and regenerating tissue. See Fig. 1A for fin region codes; numbers 0 to 2 identify regeneration stages. Circles above bars indicate significantly elevated expression (P < 0.05 in paired t-tests) in comparisons between elongated and short fin region samples (i.e., compared to the bar matching the shade of the circle).

The second step of our analysis was based on the three genes, *angptl5, cd63* and *csrp1a*, which had the strongest expression differences between the elongated medial and the short fin regions in the above analysis. Using zebrafish co-expression database, we identified eight additional candidate genes that are co-expressed with each of these genes and compared their expression levels between the caudal fin regions of *L. tigripictilis* (*arpc1, cdc42ep3, colec12, ctsk, fstl1b, mmp14b, olfml3b* and *rin3*). Among these, increased mL-expression compared to both vS- and dS-expression was detected for *colec12, ctsk, fstl1b, mmp14b* and *rin3*, whereas increased mL-expression compared to only one of the short regions (vS or dS) was detected for *arpc1 and cdc42ep3*, and no expression difference was detected for *olfml3b* (Fig. 3, Supplementary data 2).

**Figure 3:**
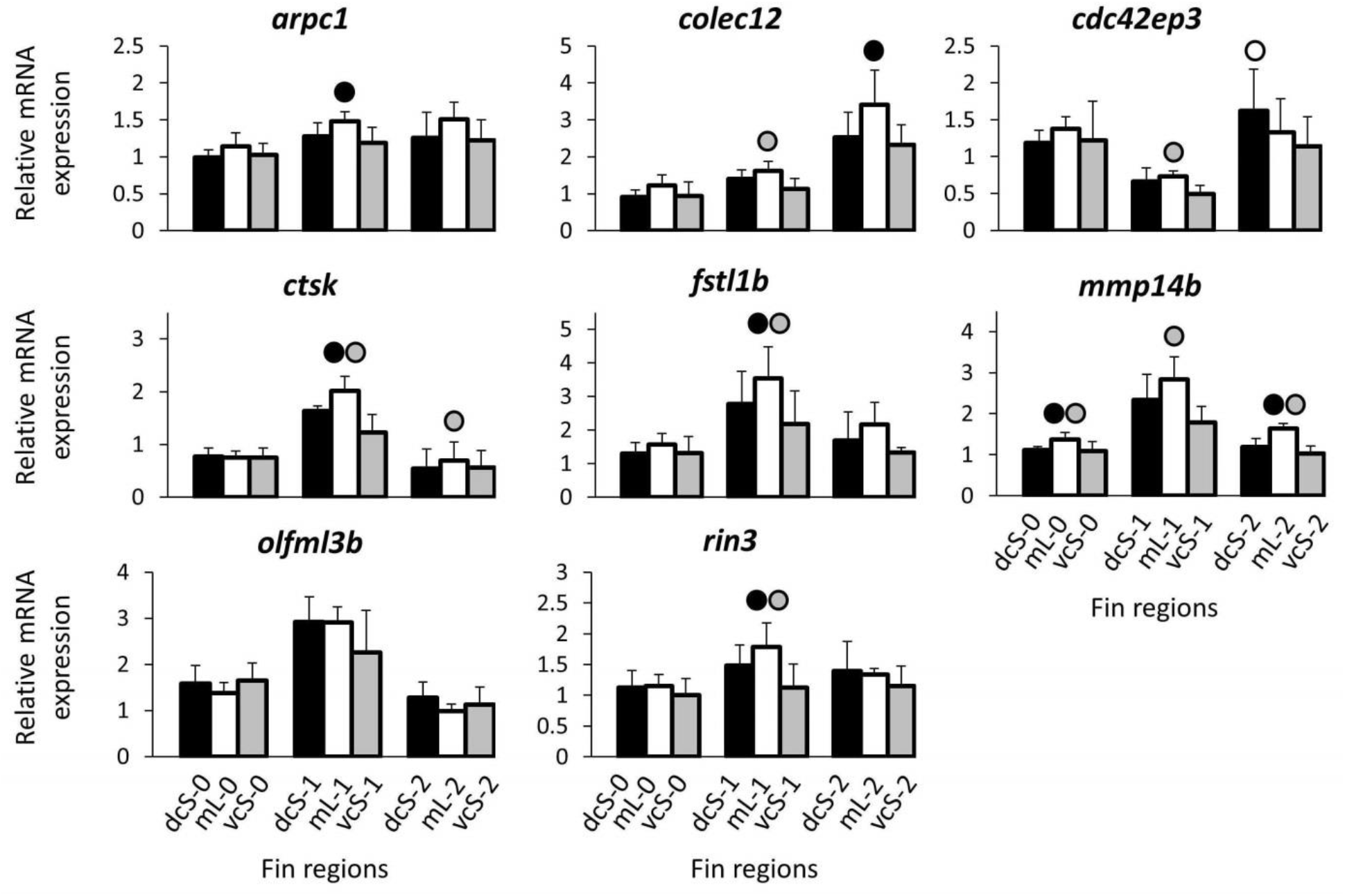
Expression levels of candidate genes selected based on co-expression with *csrp1a, angptl5* and *cd63*. Means and standard deviations of RQ in three biological replicates are shown for the elongated and short regions of the caudal in original (stage 0) and regenerating tissue. See Fig. 1A for fin region codes; numbers 0 to 2 identify regeneration stages. Circles above bars indicate significantly elevated expression (P < 0.05 in paired t-tests) in comparisons between elongated and short fin region samples (i.e., compared to the bar matching the shade of the circle).

### Expression analysis of candidate upstream regulators

Predicted upstream regulators for eight genes with the most strongly increased mL-expression in the above analyses (*angptl5, cd63, ctsk, cx43, csrp1a, fstl1b, mmp14b* and *rin3*) included the 12 transcription factors *alx4a, ap4, egr1, egr2, foxd3, foxp1, gbf1, heb, patz1, rbpj, srf*, and *znf219* (Fig. 4, Supplementary data 1). Additionally, we tested the expression of *esco2* (which was not among the predicted TFs) because it regulates *cx43* and *sema3d* in a gene regulatory network involved in the formation, growth and regeneration of fin ray segments and joints (Iovine et al., 2005; Govindan & Iovine, 2014, 2015; Banerji et al., 2016; Govindan et al., 2016). Among the 13 tested TFs, decreased mL-expression (compared to both vS- and dS-expression) was detected for *gbf1* and *rbpj* when data of all three developmental stages were pooled, but not when only stage 1 and 2 (regeneration) were analyzed. In contrast, decreased mL-expression (compared to both vS- and dS-expression) of *esco2* was detected only during regeneration. Finally, *znf219* and *alx4a* showed higher dS than vS expression, but only in analyses across all three developmental stages.

**Figure 4:**
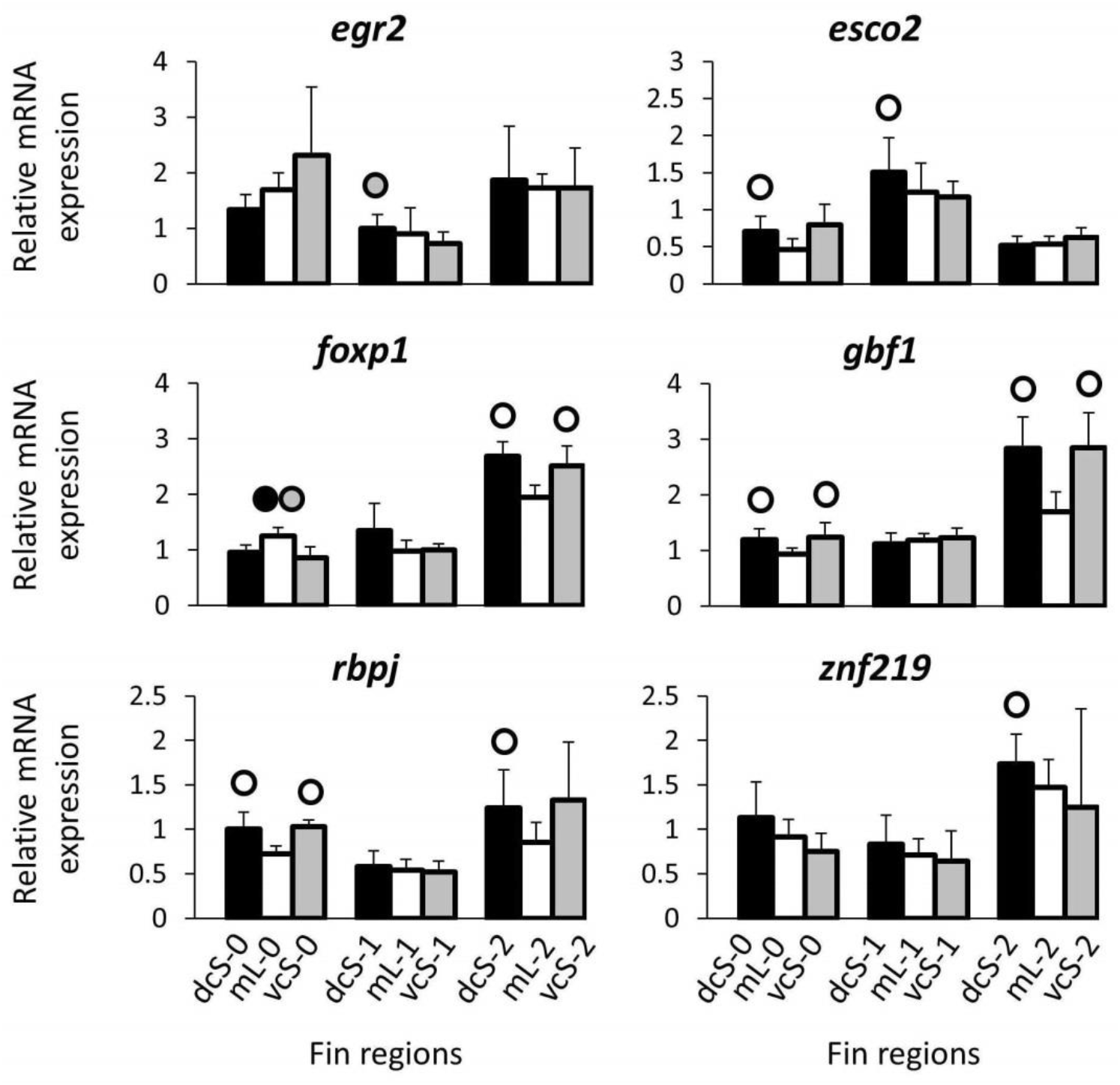
Expression levels of predicted upstream regulators. Means and standard deviations of RQ in three biological replicates are shown for the elongated and short regions of the caudal in original (stage 0) and regenerating tissue. See Fig. 1A for fin region codes; numbers 0 to 2 identify regeneration stages. Circles above bars indicate significantly elevated expression (P < 0.05 in paired t-tests) in comparisons between elongated and short fin region samples (i.e., compared to the bar matching the shade of the circle).

### Gene expression correlations

We tested for expression correlations among those of the candidate genes and upstream regulators, which had shown significant expression differences between elongated and short fin regions (Table 2). A number of significant pairwise correlations as well clusters of correlated genes were detected (Fig. 5). For instance, a cluster of positively correlated gene expression levels comprised the genes *cx43, fstl1b*, mmp14b, *c1qtnl5a, rin3, angptl5, cd63* and *ctsk*. These genes showed higher expression in the mL region in comparison to one or both of the short fin regions (dS and/or vS). The expression levels of these genes were negatively correlated with that of the transcription factor *rbpj* and positively with that of the TF *esco2*. We also detected strong positive expression correlations among the transcription factors *foxp1, gbf1* and *rbpj*. Expression of each of these TFs was negatively correlated with the expression of the TF *esco2*, and positively correlated with the expression of *pfkpa, colec12, cdc42ep3* and *dpysl5a*.

**Figure 5:**
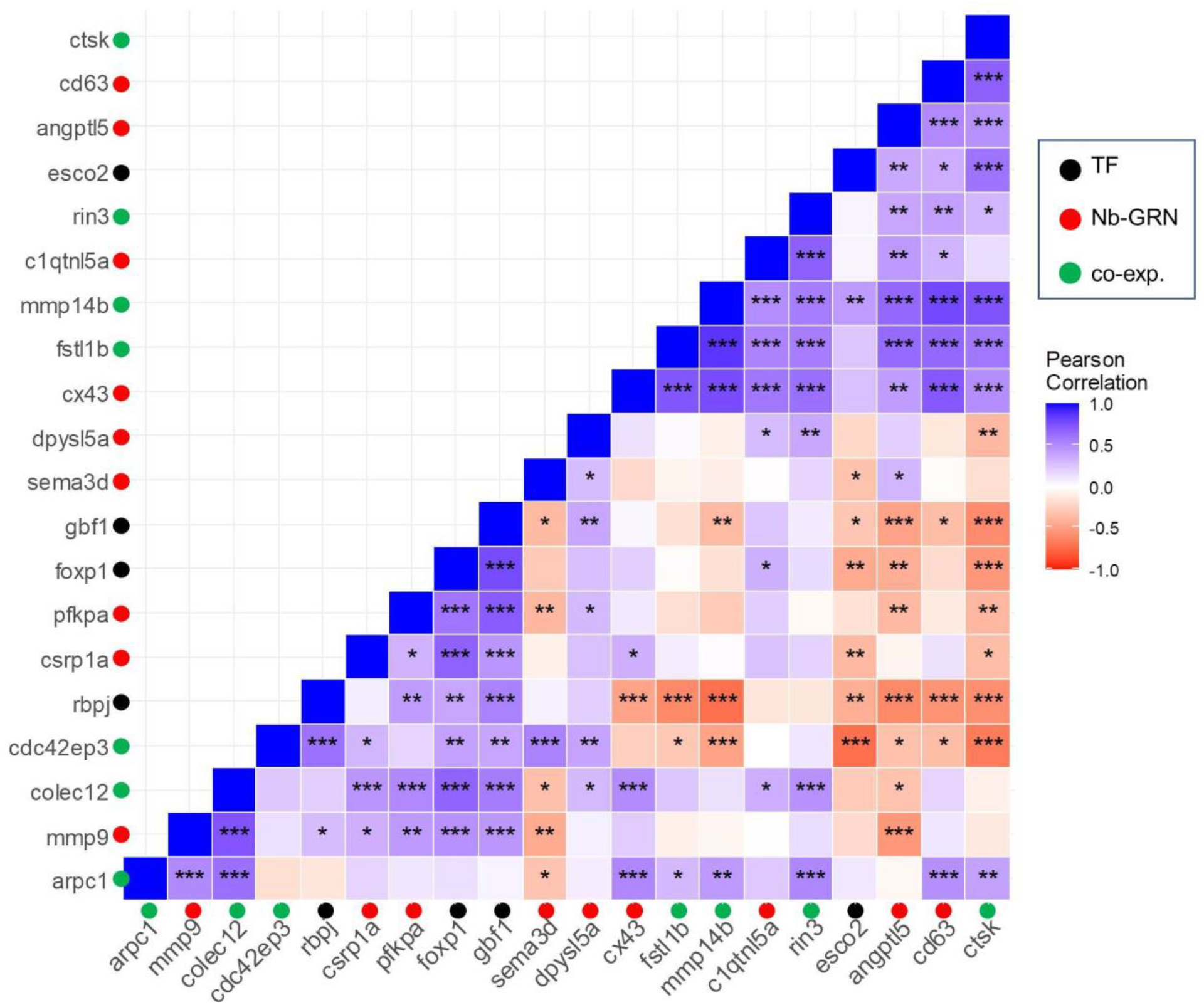
Correlation analysis reveals significant positive or negative coexpression of the candidate genes. Pearson correlation coefficient (r) was used to assess the pairwise expression similarity between the candidate genes during craniofacial development. Blue represents positive and red represents negative expression correlation. *P <0.05; **P <0.01; ***P <0.001.

## Discussion

The present study builds on previous work that revealed variation in the anatomical and molecular basis of fin shape in two phylogenetically divergent East African cichlid species (Ahi et al., 2017, 2019; Ahi & Sefc, 2018). Presently, we asked whether the mechanisms of fin elongation are more similar between two phylogenetically less divergent species, and specifically examined whether members of a GRN involved in the fin shape of *N. brichardi* might have corresponding functions in fin elongation in another lamprologine cichlid, *L. tigripictilis*. We found that both the anatomical patterns regarding the distributions of fin ray segment lengths between fin regions, as well as the molecular signals associated with the elongation of fin regions differ between the three species. As expected, similarities in molecular patterns were greater between the two Lamprologini species than in comparisons with the remotely related *S. casuarius*.

In the caudal fin of *L. tigripictilis*, the segments of the rays of the dorsal short fin region were significantly longer than the segments in both the ventral short and the medial elongated rays. This implies that the spade-like shape of the caudal fin of *L. tigripictilis* is produced by fin region-specific variation in both segment length and number. In *N. brichardi*, the ray segments in the short fin regions of the anal and dorsal fins were significantly longer than the segments in the elongated regions of the same fins (Ahi et al., 2017). Regarding the caudal fin of *N. brichardi*, we previously reported that there was no difference in segment length between the elongated and short fin regions (Ahi et al., 2017). In that analysis, segment length data of the dorsal and the ventral elongated regions of the caudal fin had been pooled into one sample representing the elongated fin rays of the caudal fin. A re-analysis of the data now revealed that the segments of the (long) dorsal rays of the caudal fin were longer than those of the (long) ventral region (mean ± SL = 289.9 ± 54.3 µm, in the dorsal region; 238.7 ± 33.0 µm, in the ventral region; N = 5 segments of 2 rays from each of 3 fish per fin region; LM, est.= 53.5, t = 2.3, p = 0.5), but in contrast to *L. tigripictilis*, the segments in the (short) medial region were equally long as those in the (long) dorsal region (mean ± SL = 291.8 ± 29.0 µm, in the medial region, N = 5 segments of 2 rays from each of 4 fish; LM, est.= 2.1, t = 0.2, p = 0.87). Segment length differences between fin regions had also been detected in *S. casuarius*, where segments were longer in the elongated than in the short regions of the anal and dorsal fins (data for the caudal fin of *S. casuarius*, which does not exhibit pronounced elongations, were not collected). Together, these data demonstrate interspecific variability regarding how segment lengths vary among elongated and short fin regions and suggest that segment lengths are not generally correlated with ray length and fin elongation in the studied cichlid species.

Gene expression differences detected between *L. tigripictilis* fin regions were partly congruent with those in *N. brichardi*. We first surveyed 16 gene members of a GRN detected in *N. brichardi*. These genes had shown significant differences in expression levels in comparisons between elongated and short fin regions in *N. brichardi*, most of them with higher expression in the elongated region. Nine of the *N*.*b*.-GRN members showed significant expression level differences between the fin regions of *L. trigripictilis*, which is a similar rate as that observed in an analogous examination of the *N*.*b*.-GRN genes in *S. casuarius* (13 out of 20 tested *N*.*b*.-GRN members with a significant signal in *S. casuarius*, Ahi et al. 2019). However, the set of genes with significant expression difference shared between *N. brichardi* and *L. tigripictilis* overlapped only partially with the genes that shared expression differences between *N. brichardi* and *S. casuarius* (Table 2).

As in the comparison between gene expression patterns in *N. brichardi* and *S. casuarius*, some but not all of the expression differences observed in *L. tigripictilis* were in the same direction as in *N. brichardi*. Among the five *N*.*b*.-GRN genes with elevated mL expression in *L. tigripictilis*, four had also shown elevated expression in the elongated regions of the fins of *N. brichardi*, and three of them also in *S. casuarius* (Table 2). The remaining members of the *N*.*b*.-GRN, for which significant expression differences were detected in *L. tigripictilis*, did not show consistent differences between each of the two elongated and the short fin regions, but rather differences between only one of the short regions and the elongated region (either dS or vS versus mL, but not both; Table 2). Among these, *mmp9* is of interest because its reduced expression in the fin region with the longest ray segments (dS in *L. tigripictilis*) is consistent with its reduced expression in the fin regions of *N. brichardi* and *S. casuarius*, which also had the longest segments (i.e., in the short regions in *N. brichardi* and in the elongated regions in *S. casuarius*). Hence, across the three species, expression of *mmp9*, which encodes a matrix remodelling enzyme with a role in fin regeneration (Yoshinari et al., 2009)(LeBert et al., 2015), is elevated in the fin regions with relatively short segments, suggesting that *mmp9* might be associated with segment length. In zebrafish, *mmp9* is negatively regulated by *cx43* (Ton & Iovine, 2013), which encodes a subunit of the gap junction protein complex and whose expression is positively correlated with segment length in the fin rays of zebrafish (Iovine et al., 2005; Sims et al., 2009). In the two lamprologine cichlid species, however, the expression levels of both genes are negatively correlated with segment length, suggesting that the interaction between *cx43* and *mmp9* and the role of *cx43* in segment growth may differ between the zebrafish and the lamprologine cichlids. Regardless of segment length, however, *cx43* was consistently more strongly expressed in the elongated fin regions in all of the studied cichlid species, which corroborates the gene as a strong candidate for a regulator of fin growth.

The elevated mL expression of *sema3d* in *L. tigripictilis* opposed the pattern in the two other cichlid species, where the gene was more strongly expressed in the short fin regions (Table 2). *Sema3d* encodes a conserved secreted ligand for several cell surface receptors involved in nervous system development, cell differentiation and bone homeostasis (Ton & Iovine, 2012; Verlinden et al., 2013), and is positively regulated by *cx43* in zebrafish (Ton & Iovine, 2012). Specifically, *sema3d* mediates the *cx43*-dependent functions in cell proliferation, joint formation and phenotypic changes of zebrafish fins (Ton & Iovine, 2012). The pattern of elevated mL expression of both *cx43* and *sema3d* in *L. tigripictilis* is therefore concordant with zebrafish and may reflect a similar functional relationship between these two genes in this species. This relationship appears, however, to be different in *N. brichardi*, where expression levels of *sema3d* and *cx43* were significantly negatively correlated. In humans, mutations in *cx43* and *sema3d* are associated with defects in finger growth (brachydactyly, (Kjaer et al., 2004; Jamsheer et al., 2014); ectrodactyly, (Sivasankaran et al., 2015), which may indicate that *cx43* and *sema3d* act in skeletal growth across much of the vertebrate tree.

In zebrafish, *cx43* is positively regulated by the expression of *esco2* (Banerji et al., 2016), and we therefore analyzed the expression levels of *esco2* in *L. tigripictilis* although the gene was not among the set of TFs predicted in the present study nor a member of the *N*.*b*.-GRN. Expression differences of *esco2* in *L. tigripictilis* were only detected in the intact fin tissue (stage 0). We did not detect the expected expression correlation between *esco2* and *cx43*, but *esco2* was significantly correlated with several other of the tested genes and TFs. Hence, data obtained for *N. brichardi* and *L. tigripictilis* (Ahi & Sefc 2018, present study) provide no evidence for a regulatory link between *esco2* and *cx43* and suggest that other transcription factors might be more important in fin morphogenesis in lamprologine cichlids. The expression correlations detected in *L. tigripictilis* suggest that elevated expression of *esco2* may negatively regulate the expression of the TFs *rbpj, gbf1* and *foxp1*, and thereby have a positive effect on several of the genes showing increased mL expression and negative expression correlations with *rbpj, gbf1* and *foxp1*.

Another gene showing consistently increased expression in elongated fin regions across the three studied cichlid species is *angptl5*, which encodes an angiopoietic protein. Its expression in endothelial cells is highly induced through interaction with osteoblasts during osteogenesis and bone remodeling in human (Simunovic et al., 2013). This indicates that increased *angptl5* expression during skeletal outgrowth is a marker for induced angiogenesis in the skeletal tissue. Increased expression of *angptl5* was also observed during exaggerated elongation of the caudal fin in swordtail fish (Kang et al., 2015).

Elevated expression in the elongated fin regions was also observed for *cd63* in both lamprologine species as well as in *S. casuarius*, and for *csrp1a* in both lamprologines, but not in *S. casuarius* (where expression levels of *csrp1a* did not differ across fin regions; Table 2). Finally, mL expression of *dpysl5a* was elevated only in comparison to vS expression during regeneration in *L. tigripictilis*. While not very strong, this pattern is in agreement with the elevated expression of *dpysl5a* in the elongated fin regions of *N. brichardi*, while no signal for *dpysl5a* was detected in *S. casuarius* (Table 2). The three genes, *cd63, csrp1a* and *dpysl5a*, are involved in neural (Monaghan et al., 2006; Ma et al., 2012; Hagedorn et al., 2016).

Given differences in gene expression patterns and inferred gene interactions between the two lamprologine species that were implied by our data, we extended our search for candidate fin shape genes in *L. tigripictilis* based on the new results. We selected the three genes with the strongest elongated-versus-short expression differences in *L. tigripictilis* and identified eight co-expressed genes in the zebrafish database. Of these, seven genes showed gene expression differences between the fin regions of *L. tigripictilis* (Table 2). These genes are not part of the GRN detected in *N. brichardi* and there are therefore no expression data for comparison with the two other cichlid species. Three of the new genes, *fstl1, colec12* and *mmp14*, have already been implicated in skeletal morphogenesis in vertebrates. Follistatin-like 1, *fstl1*, encodes a potent antagonist of the BMP pathway during skeletogenesis in vertebrates (Sylva et al., 2011) and has been shown affect digit formation in mammals (Lorda-Diez et al., 2013; Sylva et al., 2013). *Colec12* (previously known as collectin placenta protein 1 gene, *clp1*) encodes collectin-12, which is involved in vasculogenesis and has been shown to be positively associated with body elongation in zebrafish during development; i.e. knock down of *colec12* caused shortened body length in zebrafish (Fukuda et al., 2011). The metalloprotease encoding *mmp14* gene is involved in human skeletogenesis with effects on finger and toe morphology (Wilkinson et al., 2012; De Vos et al., 2018; de Vos et al., 2019). In zebrafish, mutations in *mmp14b* can lead to skeletal anomalies including shortening of body and fin, prominent frontal bone and skeletal curvatures (De Vos et al., 2018). The elevated expression levels of *mmp14* and *colec12* in elongated compared to short fin regions in *L. tigripictilis* are consistent with the mutant zebrafish phenotypes.

We also extended our candidate gene set by predicting transcription factors for genes with strongly increased mL-expression in *L. tigripictilis*. The 12 predicted TFs included three that had already been proposed as upstream regulators of the GRN in *N. brichardi* (*egr2* and *foxd3* with elevated expression in the elongated fin regions, and *foxp1* with elevated expression in the short fin regions; (Ahi & Sefc 2018). We detected no significant expression differences among fin regions for *egr2* and *foxd3* in *L. tigripictilis* and only a weak signal for *foxp1*, which matched the pattern in *N. brichardi* (Table 2). This may suggest transcriptional repressor activity of foxp1 which is also implicated as the main potential repressor TF involved in formation of lip hypertrophy (Lecaudey et al., 2021) and exaggerated snout in two other East African cichlid species. The absence of significant expression differences in two of the proposed upstream regulators of the *N*.*b*.- GRN further corroborates the supposition that there is only limited overlap in the molecular mechanisms of fin shape formation between the two lamprologine cichlids.

TFs predicted from genes with strongly increased mL-expression are expected to display expression correlations with these genes and therefore to display either increased or decreased mL expression. The expected pattern was observed for *gbf1* and *rbpj*, both of which showed reduced mL expression compared to both dS and vS fin regions. The expression levels of *gbf1* and *rbpj* were also significantly correlated with the majority of the genes involved in the prediction procedure (Fig. 5). Golgi brefeldin A-resistant factor 1 gene, *gbf1*, encodes a protein that functions as a guanine nucleotide exchange factor and plays important roles in regulating organelle structure and cargo-selective vesicle trafficking (Manolea et al., 2008). During zebrafish development, *gbf1* is involved in vascular system formation, pigmentation and morphogenesis of the caudal fin (Chen et al., 2017). Knockdown of *GBF1* in mammalian cells leads to a range of structural anomalies which eventually inhibits trafficking of transmembrane proteins and cell death (Citterio et al., 2008).

The second TF, *rbpj*, encodes a transcription factor with dual regulatory activities, which can act as a an activator or repressor depending on its interaction with Notch signal proteins (Castel et al., 2013). For instance, the Notch intracellular domain (NICD) and *rbpj* form a complex that can act as transcription repressor and negatively regulate chondrocyte differentiation (Chen et al., 2013). On the other hand, the NICD-rbpj complex acts as transcriptional activator inducing osteoblast proliferation (Tao et al., 2010). Moreover, *rbpj* has been shown to inhibit osteoclastogenesis and bone resorption (Zhao et al., 2012; Miller et al., 2016). It should be noted that Notch signal activity and *rbpj* transcription are both required for maintaining blastema cells in a plastic, undifferentiated and proliferative state, which is essential for fin regeneration in zebrafish (Münch et al., 2013). In human, mutation in *Rbpj* is shown to be associated with etiology of Adams-Oliver syndrome (AOS) which is identified with multiple-malformation disorders, and particularly, with terminal limb defects (Hassed et al., 2012; Nakayama et al., 2014). The terminal limb defects in AOS are characterized by shortening of the end of the fingers (brachydactyly) and curvature of the digits (clinodactyily) (Nakayama et al., 2014). In mammals, *rbpj* can act as upstream regulator of *mmp14* expression (Gao et al., 2016; Nus et al., 2016). The two genes show opposite expression patterns (*mmp14* with elevated and *rbpj* with reduced mL expression, respectively) and significantly negative expression correlations (r = -0.72, p > 0.001) in *L. tigripictilis*, suggesting that a regulatory interaction also exists in this cichlid species.

## Conclusions

Our data suggest that the elongation of the medial region of the caudal fin of *L. tigripictilis*, which causes the spade shape outline of the fin, emanates from different gene regulatory mechanisms than those implemented in fin region elongation in the related species *N. brichardi*. Only few of the GRN members, that were proposed to be involved in the filamentous elongation of fin edges in *N. brichardi*, also showed the expected, elevated expression in the elongated region of the *L. tigripictilis* fin. Moreover, the predicted upstream regulators of the genes, whose expression was statistically associated with elongated fin regions, were also different in *L. tigripictilis* (*gbf1* and *rbpj*) from those found in *N. brichardi*. This raises the possibility of diverse and distinct molecular processes underling fin morphogenesis even within a fish taxon. Alternatively, the incongruences between the transcriptional patterns of the two species may also represent positional effects, i.e. the fin elongation in the medial region of the fin may involve different genes than fin elongation at dorsal or ventral edges. Strikingly, we also found that the molecular mechanism defining fin segment length (which probably involves the *esco2* transcription factor) is decoupled from the mechanism determining the overall fin length. In other words, a mechanism determining segment numbers but not segment lengths might be the key player in medial fin elongation in *L. tigripictilis*. Collated with previous results, our data show substantial diversity in the regulation of fin shape among cichlid fishes and also in comparison to zebrafish.

## Declarations

### Authors’ contributions

EPA and KMS designed the study, analysed the expression data and wrote the manuscript. EPA conducted the fin dissections, RNA extraction and qPCR laboratory experiments. EPA and KMS prepared figures for gene expression sections. FR conducted fin skeletal staining, photography, measurement and analysis of fin ray segments as well as the schematic drawing of fish (Fig. 1A and B).

## Acknowledgements

The authors thank Wolfgang Gessl (www.pisces.at) for his responsible management of our fish. We also thank Holger Zimmermann and Stephan Koblmüller for sharing their precious knowledge on cichlid fishes of Lake Tanganyika, and Martin Grube and his lab for technical assistance and access to their real-time PCR System. The authors acknowledge the financial support by the University of Graz.

## Availability of data and materials

All the gene expression data generated or analysed during this study are included in this published article.

## Ethics approval and consent to participate

All experimental protocols were approved by the Federal Ministry of Science, Research and Economy of Austria (BMWFW) under permit BMWFW-66.007/0004-WF/V/3b/2016. All methods were carried out in accordance with relevant guidelines and regulations of the Austrian animal welfare law. All methods are reported in accordance with ARRIVE guidelines (https://arriveguidelines.org) for the reporting of animal experiments.

---

**Supplementary data 1. Primer information, selection of co-expressed genes and TF binding site enrichments**.

**Supplementary data 2. Statistical information**.

## References

Ahi, E. P., J. Guðbrandsson, K. H. Kapralova, S. R. Franzdóttir, S. S. Snorrason, V. H. Maier, & Z. O. Jónsson, 2013. Validation of Reference Genes for Expression Studies during Craniofacial Development in Arctic Charr. PloS one 8: e66389.

Ahi, E. P., L. A. Lecaudey, A. Ziegelbecker, O. Steiner, W. Goessler, & K. M. Sefc, 2020. Expression levels of the tetratricopeptide repeat protein gene ttc39b covary with carotenoid-based skin colour in cichlid fish. Biology Letters.

Ahi, E. P., F. Richter, L. A. Lecaudey, & K. M. Sefc, 2019. Gene expression profiling suggests differences in molecular mechanisms of fin elongation between cichlid species. Scientific Reports Nature Publishing Group 9:.

Ahi, E. P., F. Richter, & K. M. Sefc, 2017. A gene expression study of ornamental fin shape in Neolamprologus brichardi, an African cichlid species. Scientific Reports.

Ahi, E. P., & K. M. Sefc, 2018. Towards a gene regulatory network shaping the fins of the Princess cichlid. Scientific Reports Nature Publishing Group 8: 9602.

Ahi, E. P., S. S. Steinhäuser, A. Pálsson, S. R. Franzdóttir, S. S. Snorrason, V. H. Maier, & Z. O. Jónsson, 2015. Differential expression of the aryl hydrocarbon receptor pathway associates with craniofacial polymorphism in sympatric Arctic charr. EvoDevo 6:.

Andersen, C. L., J. L. Jensen, & T. F. Ørntoft, 2004. Normalization of real-time quantitative reverse transcription-PCR data: a model-based variance estimation approach to identify genes suited for normalization, applied to bladder and colon cancer data sets. Cancer research 64: 5245–5250, http://www.ncbi.nlm.nih.gov/pubmed/15289330.

Bailey, T. L., M. Boden, F. A. Buske, M. Frith, C. E. Grant, L. Clementi, J. Ren, W. W. Li, & W. S. Noble, 2009. MEME SUITE: tools for motif discovery and searching. Nucleic acids research Oxford University Press 37: W202–8.

Banerji, R., D. M. Eble, M. K. Iovine, & R. V Skibbens, 2016. Esco2 regulates cx43 expression during skeletal regeneration in the zebrafish fin. Developmental dynamics : an official publication of the American Association of Anatomists 245: 7–21.

Benjamini, Y., & Y. Hochberg, 1995. Controlling the false discovery rate: A Practical and powerful approach to multiple testing. J.Roy.Statist.Soc. 57: 289–300.

Brawand, D., C. E. Wagner, Y. I. Li, M. Malinsky, I. Keller, S. Fan, O. Simakov, A. Y. Ng, Z. W. Lim, E. Bezault, J. Turner-Maier, J. Johnson, R. Alcazar, H. J. Noh, P. Russell, B. Aken, J. Alföldi, C. Amemiya, N. Azzouzi, J.-F. Baroiller, F. Barloy-Hubler, A. Berlin, R. Bloomquist, K. L. Carleton, M. a. Conte, H. D’Cotta, O. Eshel, L. Gaffney, F. Galibert, H. F. Gante, S. Gnerre, L. Greuter, R. Guyon, N. S. Haddad, W. Haerty, R. M. Harris, H. a. Hofmann, T. Hourlier, G. Hulata, D. B. Jaffe, M. Lara, A. P. Lee, I. MacCallum, S. Mwaiko, M. Nikaido, H. Nishihara, C. Ozouf-Costaz, D. J. Penman, D. Przybylski, M. Rakotomanga, S. C. P. Renn, F. J. Ribeiro, M. Ron, W. Salzburger, L. Sanchez-Pulido, M. E. Santos, S. Searle, T. Sharpe, R. Swofford, F. J. Tan, L. Williams, S. Young, S. Yin, N. Okada, T. D. Kocher, E. a. Miska, E. S. Lander, B. Venkatesh, R. D. Fernald, A. Meyer, C. P. Ponting, J. T. Streelman, K. Lindblad-Toh, O. Seehausen, & F. Di Palma, 2014. The genomic substrate for adaptive radiation in African cichlid fish. Nature 513: 375–381.

Castel, D., P. Mourikis, S. J. J. Bartels, A. B. Brinkman, S. Tajbakhsh, & H. G. Stunnenberg, 2013. Dynamic binding of RBPJ is determined by notch signaling status. Genes and Development Cold Spring Harbor Laboratory Press 27: 1059–1071.

Chen, J., X. Wu, L. Yao, L. Yan, L. Zhang, J. Qiu, X. Liu, S. Jia, & A. Meng, 2017. Impairment of cargo transportation caused by gbf1 mutation disrupts vascular integrity and causes hemorrhage in zebrafish embryos. Journal of Biological Chemistry American Society for Biochemistry and Molecular Biology Inc. 292: 2315–2327.

Chen, S., J. Tao, Y. Bae, M.-M. Jiang, T. Bertin, Y. Chen, T. Yang, & B. Lee, 2013. Notch gain of function inhibits chondrocyte differentiation via Rbpj-dependent suppression of Sox9. Journal of Bone and Mineral Research 28: 649–659, http://www.pubmedcentral.nih.gov/articlerender.fcgi?artid=3548081&tool=pmcentrez&rendertype=abstract.

Christen, B., V. Robles, M. Raya, I. Paramonov, & J. C. Izpisúa Belmonte, 2010. Regeneration and reprogramming compared. BMC biology BioMed Central 8: 5.

Citterio, C., A. Vichi, G. Pacheco-Rodriguez, A. M. Aponte, J. Moss, & M. Vaughan, 2008. Unfolded protein response and cell death after depletion of brefeldin A-inhibited guanine nucleotide-exchange protein GBF. Proceedings of the National Academy of Sciences of the United States of America National Academy of Sciences 105: 2877–2882.

De Vos, I. J. H. M., E. Y. Tao, S. L. M. Ong, J. L. Goggi, T. Scerri, G. R. Wilson, C. G. M. Low, A. S. W. Wong, D. Grussu, A. P. A. Stegmann, M. Van Geel, R. Janssen, D. J. Amor, M. Bahlo, N. R. Dunn, T. J. Carney, P. J. Lockhart, B. J. Coull, & M. A. M. Van Steensel, 2018. Functional analysis of a hypomorphic allele shows that MMP14 catalytic activity is the prime determinant of the Winchester syndrome phenotype. Human Molecular Genetics Oxford University Press 27: 2775–2788.

de Vos, I. J. H. M., A. S. W. Wong, T. J. M. Welting, B. J. Coull, & M. A. M. van Steensel, 2019. Multicentric osteolytic syndromes represent a phenotypic spectrum defined by defective collagen remodeling. American Journal of Medical Genetics, Part A. Wiley-Liss Inc., 1652–1664.

Flicek, P., I. Ahmed, M. R. Amode, D. Barrell, K. Beal, S. Brent, D. Carvalho-Silva, P. Clapham, G. Coates, S. Fairley, S. Fitzgerald, L. Gil, C. García-Girón, L. Gordon, T. Hourlier, S. Hunt, T. Juettemann, A. K. Kähäri, S. Keenan, M. Komorowska, E. Kulesha, I. Longden, T. Maurel, W. M. McLaren, M. Muffato, R. Nag, B. Overduin, M. Pignatelli, B. Pritchard, E. Pritchard, H. S. Riat, G. R. S. Ritchie, M. Ruffier, M. Schuster, D. Sheppard, D. Sobral, K. Taylor, A. Thormann, S. Trevanion, S. White, S. P. Wilder, B. L. Aken, E. Birney, F. Cunningham, I. Dunham, J. Harrow, J. Herrero, T. J. P. Hubbard, N. Johnson, R. Kinsella, A. Parker, G. Spudich, A. Yates, A. Zadissa, & S. M. J. Searle, 2013. Ensembl 2013. Nucleic acids research Oxford University Press 41: D48–55.

Fukuda, M., K. Ohtani, S. J. Jang, T. Yoshizaki, K. I. Mori, W. Motomura, I. Yoshida, Y. Suzuki, Y. Kohgo, & N. Wakamiya, 2011. Molecular cloning and functional analysis of scavenger receptor zebrafish CL-P1. Biochimica et Biophysica Acta - General Subjects Elsevier 1810: 1150–1159.

Gao, X., D. Han, & W. Fan, 2016. Down-regulation of RBP-J mediated by microRNA-133a suppresses dendritic cells and functions as a potential tumor suppressor in osteosarcoma. Experimental Cell Research Academic Press Inc. 349: 264–272.

Govindan, J., & M. K. Iovine, 2014. Hapln1a Is Required for Connexin43-Dependent Growth and Patterning in the Regenerating Fin Skeleton. PLoS ONE Public Library of Science 9: e88574.

Govindan, J., & M. K. Iovine, 2015. Dynamic remodeling of the extra cellular matrix during zebrafish fin regeneration. Gene Expression Patterns 19: 21–29.

Govindan, J., K. M. Tun, & M. K. Iovine, 2016. Cx43-Dependent Skeletal Phenotypes Are Mediated by Interactions between the Hapln1a-ECM and Sema3d during Fin Regeneration. PLOS ONE Public Library of Science 11: e0148202.

Hagedorn, M., G. Siegfried, K. B. Hooks, & A.-M. Khatib, 2016. Integration of zebrafish fin regeneration genes with expression data of human tumors in silico uncovers potential novel melanoma markers. Oncotarget Impact Journals, LLC 7: 71567–71579.

Hasegawa, T., C. J. Hall, P. S. Crosier, G. Abe, K. Kawakami, A. Kudo, & A. Kawakami, 2017. Transient inflammatory response mediated by interleukin-1β is required for proper regeneration in zebrafish fin fold. eLife eLife Sciences Publications, Ltd 6: e22716.

Hassed, S. J., G. B. Wiley, S. Wang, J. Y. Lee, S. Li, W. Xu, Z. J. Zhao, J. J. Mulvihill, J. Robertson, J. Warner, & P. M. Gaffney, 2012. RBPJ mutations identified in two families affected by Adams-Oliver syndrome. American Journal of Human Genetics Cell Press 91: 391–395.

Hellemans, J., G. Mortier, A. De Paepe, F. Speleman, & J. Vandesompele, 2007. qBase relative quantification framework and software for management and automated analysis of real-time quantitative PCR data. Genome biology 8: R19, http://www.pubmedcentral.nih.gov/articlerender.fcgi?artid=1852402&tool=pmcentrez&rendertype=abstract.

Huang, J., & L. Chen, 2017. IL-1β inhibits osteogenesis of human bone marrow-derived mesenchymal stem cells by activating FoxD3/microRNA-496 to repress wnt signaling. genesis e23040.

Iovine, M. K., 2007. Conserved mechanisms regulate outgrowth in zebrafish fins. Nature Chemical Biology. Nature Publishing Group, 613–618.

Iovine, M. K., E. P. Higgins, A. Hindes, B. Coblitz, & S. L. Johnson, 2005. Mutations in connexin43 (GJA1) perturb bone growth in zebrafish fins. Developmental Biology 278: 208–219.

Irisarri, I., P. Singh, S. Koblmüller, J. Torres-Dowdall, F. Henning, P. Franchini, C. Fischer, A. R. Lemmon, E. M. Lemmon, G. G. Thallinger, C. Sturmbauer, & A. Meyer, 2018. Phylogenomics uncovers early hybridization and adaptive loci shaping the radiation of Lake Tanganyika cichlid fishes. Nature Communications Nature Publishing Group 9: 1–12.

Jamsheer, A., A. Sowińska-Seidler, M. Socha, A. Stembalska, C. Kiraly-Borri, & A. Latos-Bieleńska, 2014. Three novel GJA1 missense substitutions resulting in oculo-dento-digital dysplasia (ODDD) - Further extension of the mutational spectrum. Gene Elsevier 539: 157–161.

Kang, J. H., T. Manousaki, P. Franchini, S. Kneitz, M. Schartl, & A. Meyer, 2015. Transcriptomics of two evolutionary novelties: how to make a sperm-transfer organ out of an anal fin and a sexually selected “sword” out of a caudal fin. Ecology and Evolution 5: 848–864.

Kjaer, K. W., L. Hansen, H. Eiberg, P. Leicht, J. M. Opitz, & N. Tommerup, 2004. Novel Connexin 43 (GJA1) mutation causes oculo-dento-digital dysplasia with curly hair. American Journal of Medical Genetics Wiley-Liss Inc. 127A: 152–157.

LeBert, D. C., J. M. Squirrell, J. Rindy, E. Broadbridge, Y. Lui, A. Zakrzewska, K. W. Eliceiri, A. H. Meijer, & A. Huttenlocher, 2015. Matrix metalloproteinase 9 modulates collagen matrices and wound repair. Development (Cambridge) Company of Biologists Ltd 142: 2136–2146.

Lecaudey, L. A., P. Singh, C. Sturmbauer, A. Duenser, W. Gessl, & E. P. Ahi, 2021. Transcriptomics unravels molecular players shaping dorsal lip hypertrophy in the vacuum cleaner cichlid, Gnathochromis permaxillaris. BMC genomics NLM (Medline) 22: 506.

Lecaudey, L. A., C. Sturmbauer, P. Singh, & E. P. Ahi, 2019. Molecular mechanisms underlying nuchal hump formation in dolphin cichlid, Cyrtocara moorii. Scientific Reports 9: 20296, http://www.nature.com/articles/s41598-019-56771-7.

Li, L., B. Yan, Y.-Q. Shi, W.-Q. Zhang, & Z.-L. Wen, 2012. Live imaging reveals differing roles of macrophages and neutrophils during zebrafish tail fin regeneration. The Journal of biological chemistry American Society for Biochemistry and Molecular Biology 287: 25353–25360.

Lorda-Diez, C. I., J. A. Montero, J. Rodriguez-Leon, J. A. Garcia-Porrero, & J. M. Hurle, 2013. Expression and Functional Study of Extracellular BMP Antagonists during the Morphogenesis of the Digits and Their Associated Connective Tissues. PLoS ONE 8: e60423–e60423.

Luehr, S., H. Hartmann, & J. Söding, 2012. The XXmotif web server for eXhaustive, weight matriX-based motif discovery in nucleotide sequences. Nucleic acids research Oxford University Press 40: W104–9.

Ma, L., Y.-M. Yu, Y. Guo, R. P. Hart, & M. Schachner, 2012. Cysteine- and glycine-rich protein 1a is involved in spinal cord regeneration in adult zebrafish. The European journal of neuroscience NIH Public Access 35: 353–365.

Mahony, S., & P. V Benos, 2007. STAMP: a web tool for exploring DNA-binding motif similarities. Nucleic acids research 35: W253–8, http://www.pubmedcentral.nih.gov/articlerender.fcgi?artid=1933206&tool=pmcentrez&rendertype=abstract.

Manolea, F., A. Claude, J. Chun, J. Rosas, & P. Melançon, 2008. Distinct functions for Arf guanine nucleotide exchange factors at the Golgi complex: GBF1 and BIGs are required for assembly and maintenance of the Golgi stack and trans-Golgi network, respectively. Molecular Biology of the Cell American Society for Cell Biology 19: 523–535.

Matys, V., E. Fricke, R. Geffers, E. Gössling, M. Haubrock, R. Hehl, K. Hornischer, D. Karas, A. E. Kel, O. V Kel-Margoulis, D.-U. Kloos, S. Land, B. Lewicki-Potapov, H. Michael, R. Münch, I. Reuter, S. Rotert, H. Saxel, M. Scheer, S. Thiele, & E. Wingender, 2003. TRANSFAC: transcriptional regulation, from patterns to profiles. Nucleic acids research Oxford University Press 31: 374–378.

Miller, C. H., S. M. Smith, M. Elguindy, T. Zhang, J. Z. Xiang, X. Hu, L. B. Ivashkiv, & B. Zhao, 2016. RBP-J–Regulated miR-182 Promotes TNF-α–Induced Osteoclastogenesis. The Journal of Immunology The American Association of Immunologists 196: 4977–4986.

Monaghan, J. R., J. A. Walker, R. B. Page, S. Putta, C. K. Beachy, & S. R. Voss, 2006. Early gene expression during natural spinal cord regeneration in the salamander Ambystoma mexicanum. Journal of Neurochemistry Blackwell Publishing Ltd 101: 27–40.

Münch, J., A. González-Rajal, & J. L. de la Pompa, 2013. Notch regulates blastema proliferation and prevents differentiation during adult zebrafish fin regeneration. Development (Cambridge, England) 140: 1402–1411.

Nakatani, Y., M. Nishidate, M. Fujita, A. Kawakami, & A. Kudo, 2007. Migration of mesenchymal cell fated to blastema is necessary for fish fin regeneration. Development, Growth & Differentiation Blackwell Publishing Asia 50: 71–83.

Nakayama, T., H. Saitsu, W. Endo, A. Kikuchi, M. Uematsu, K. Haginoya, N. Hino-fukuyo, T. Kobayashi, M. Iwasaki, T. Tominaga, S. Kure, & N. Matsumoto, 2014. RBPJ is disrupted in a case of proximal 4p deletion syndrome with epilepsy. Brain and Development Elsevier 36: 532–536.

Nus, M., B. Martinez-Poveda, D. MacGrogan, R. Chevre, G. D’Amato, M. Sbroggio, C. Rodriguez, J. Martinez-Gonzalez, V. Andrés, A. Hidalgo, & J. L. De La Pompa, 2016. Endothelial Jag1-RBPJ signalling promotes inflammatory leucocyte recruitment and atherosclerosis. Cardiovascular Research Oxford University Press 112: 568–580.

Obayashi, T., & K. Kinoshita, 2011. COXPRESdb: a database to compare gene coexpression in seven model animals. Nucleic acids research 39: D1016–22, http://www.pubmedcentral.nih.gov/articlerender.fcgi?artid=3013720&tool=pmcentrez&rendertype=abstract.

Pfaffl, M. W., 2001. A new mathematical model for relative quantification in real-time RT-PCR. Nucleic acids research 29: e45, http://www.pubmedcentral.nih.gov/articlerender.fcgi?artid=55695&tool=pmcentrez&rendertype=abstract.

Pfaffl, M. W., A. Tichopad, C. Prgomet, & T. P. Neuvians, 2004. Determination of stable housekeeping genes, differentially regulated target genes and sample integrity: BestKeeper--Excel-based tool using pair-wise correlations. Biotechnology letters 26: 509–515, http://www.ncbi.nlm.nih.gov/pubmed/15127793.

Pfefferli, C., & A. Jaźwińska, 2015. The art of fin regeneration in zebrafish. Regeneration Wiley 2: 72–83, https://onlinelibrary.wiley.com/doi/10.1002/reg2.33.

Rabinowitz, J. S., A. M. Robitaille, Y. Wang, C. A. Ray, R. Thummel, H. Gu, D. Djukovic, D. Raftery, J. D. Berndt, & R. T. Moon, 2017. Transcriptomic, proteomic, and metabolomic landscape of positional memory in the caudal fin of zebrafish. Proceedings of the National Academy of Sciences of the United States of America National Academy of Sciences 114: E717–E726.

Ramakers, C., J. M. Ruijter, R. H. L. Deprez, & A. F. M. Moorman, 2003. Assumption-free analysis of quantitative real-time polymerase chain reaction (PCR) data. Neuroscience letters 339: 62–66, http://www.ncbi.nlm.nih.gov/pubmed/12618301.

Santos, M. E., L. Baldo, L. Gu, N. Boileau, Z. Musilova, & W. Salzburger, 2016. Comparative transcriptomics of anal fin pigmentation patterns in cichlid fishes. BMC Genomics BioMed Central 17: 712.

Sehring, I. M., & G. Weidinger, 2020. Recent advancements in understanding fin regeneration in zebrafish. WIREs Developmental Biology John Wiley and Sons Inc. 9: e367, https://onlinelibrary.wiley.com/doi/10.1002/wdev.367.

Sims, K., D. M. Eble, & M. K. Iovine, 2009. Connexin43 regulates joint location in zebrafish fins. Developmental Biology 327: 410–418.

Simunovic, F., D. Steiner, D. Pfeifer, G. B. Stark, G. Finkenzeller, & F. Lampert, 2013. Increased extracellular matrix and proangiogenic factor transcription in endothelial cells after cocultivation with primary human osteoblasts. Journal of Cellular Biochemistry John Wiley & Sons, Ltd 114: 1584–1594.

Singh, P., C. Börger, H. More, & C. Sturmbauer, 2017. The role of alternative splicing and differential gene expression in cichlid adaptive radiation. Genome Biology and Evolution 9: 2764–2781.

Sivasankaran, A., A. Srikanth, P. S. Kulshreshtha, D. Anuradha, J. S. Kadandale, & C. R. Samuel, 2015. Split Hand/Foot Malformation Associated with 7q21.3 Microdeletion: A Case Report. Molecular Syndromology S. Karger AG 6: 287–296.

Sylva, M., V. S. W. Li, A. A. A. Buffing, J. H. van Es, M. van den Born, S. van der Velden, Q. Gunst, J. H. Koolstra, A. F. M. Moorman, H. Clevers, & M. J. B. van den Hoff, 2011. The BMP Antagonist follistatin-like 1 is required for skeletal and lung organogenesis. PLoS ONE Public Library of Science 6:.

Sylva, M., A. F. M. Moorman, & M. J. B. van den Hoff, 2013. Follistatin-like 1 in vertebrate development. Birth Defects Research Part C: Embryo Today: Reviews John Wiley & Sons, Ltd 99: 61–69.

Tao, J., S. Chen, T. Yang, B. Dawson, E. Munivez, T. Bertin, & B. Lee, 2010. Osteosclerosis owing to Notch gain of function is solely Rbpj-dependent. Journal of Bone and Mineral Research John Wiley & Sons, Ltd 25: 2175–2183.

Ton, Q. V., & K. M. Iovine, 2012. Semaphorin3d mediates Cx43-dependent phenotypes during fin regeneration. Developmental Biology 366: 195–203.

Ton, Q. V., & M. K. Iovine, 2013. Identification of an evx1-Dependent Joint-Formation Pathway during FIN Regeneration. PLoS ONE Public Library of Science 8: e81240.

Vandesompele, J., K. De Preter, F. Pattyn, B. Poppe, N. Van Roy, A. De Paepe, & F. Speleman, 2002. Accurate normalization of real-time quantitative RT-PCR data by geometric averaging of multiple internal control genes. Genome biology 3: RESEARCH0034, http://www.pubmedcentral.nih.gov/articlerender.fcgi?artid=126239&tool=pmcentrez&rendertype=abstract.

Verlinden, L., C. Kriebitzsch, I. Beullens, B. K. Tan, G. Carmeliet, & A. Verstuyf, 2013. Nrp2 deficiency leads to trabecular bone loss and is accompanied by enhanced osteoclast and reduced osteoblast numbers. Bone 55: 465–475.

Walker, M., & C. Kimmel, 2007. A two-color acid-free cartilage and bone stain for zebrafish larvae. Biotechnic & Histochemistry 82: 23–28.

Wehner, D., & G. Weidinger, 2015. Signaling networks organizing regenerative growth of the zebrafish fin. Trends in Genetics. Elsevier Ltd, 336–343.

Wilkinson, J. M., R. K. Davidson, T. E. Swingler, E. R. Jones, A. N. Corps, P. Johnston, G. P. Riley, A. J. Chojnowski, & I. M. Clark, 2012. MMP-14 and MMP-2 are key metalloproteases in Dupuytren’s disease fibroblast-mediated contraction. Biochimica et Biophysica Acta - Molecular Basis of Disease Elsevier 1822: 897–905.

Yoshinari, N., T. Ishida, A. Kudo, & A. Kawakami, 2009. Gene expression and functional analysis of zebrafish larval fin fold regeneration. Developmental Biology Academic Press Inc. 325: 71–81.

Zhao, B., S. N. Grimes, S. Li, X. Hu, & L. B. Ivashkiv, 2012. TNF-induced osteoclastogenesis and inflammatory bone resorption are inhibited by transcription factor RBP-J. Journal of Experimental Medicine The Rockefeller University Press 209: 319–334.

